# A multifaceted approach for analyzing complex phenotypic data in rodent models of autism

**DOI:** 10.1101/439455

**Authors:** Ishita Das, Marcel A. Estevez, Anjali A. Sarkar, Sharmila Banerjee-Basu

## Abstract

AutDB features a modular framework that aims at collating multifactorial risk factors associated with autism spectrum disorder (ASD). The animal model (AM) module of AutDB was first developed for mouse models of genes and CNVs associated with ASD (Kumar et al., 2011). Subsequently, environmentally induced rodent models were introduced to capture the full spectrum of risk-factors associated with ASD, along with idiopathic models represented by inbred strains. Using the data systematically annotated in AutDB, we depict the intricate trends in the research findings based on rodent models of ASD. We identify the top 30 most frequently studied phenotypes extracted from 911 genetic, 269 induced and 17 inbred rodent models of ASD extracted from 787 publications. As expected, many of these include animal model equivalents of the ‘core’ phenotypes associated with ASD, as well as several comorbid features of ASD including anxiety, seizures and motor-control deficits. Uniquely, AutDB curates rescue models where various treatment strategies were used in rodent ASD models to alleviate ASD-relevant phenotypes. We further examine ASD models based on 52 genes and 2 CNV loci to identify 24 pharmaceutical agents that were used in 2 or more paradigms for testing their efficacy. As a case study, we analyze various Shank3 mouse models providing a high-resolution view of the *in vivo* role of this high-confidence ASD gene. Together, this resource provides a snapshot of genetic and induced models of ASD within a shared annotation platform to examine the complex meshing of diverse ASD-associated risk-factors.

## Introduction

Animal models have been pivotal in understanding the etiology of many human diseases and determining effective therapeutic interventions (Smithies, 1993). Research using animal models has unearthed mechanistic underpinnings and identified therapeutic targets for neurological disorders arising due to dysfunction of specific cell-types or brain regions, e.g., Parkinson’s disease (Zeng et al., 2018). Rodent models for diseases caused by viral or bacterial infections including some types of cancer and acquired immune deficiency syndrome (AIDS) have also led to an understanding of the fundamentals of the mammalian immune system leading to practical advances in healthcare management (Curwen and Wedge, 2009; von Bubnoff, 2008; Washington et al., 2013). However, for complex behavioral disorders that have a more diffused pattern, with multiple and sporadic genomic loci implicated in disease development, rodent models have evoked more questions than answers, be it schizophrenia, Down syndrome or ASD (Chadman et al., 2012; O’Donnell, 2013; Victorino et al., 2017).

AutDB has focused on curating and annotating ASD research for the past 10 years using a scientific annotation framework rooted in the biology of the disorder (Basu et al., 2009) The systematic annotation of autism-related data on a standardized platform has been an invaluable resource to researchers seeking to sort through confounding and ground-breaking findings. Toward this end, ASD-associated AutDB gene and CNV datasets have been used widely by the research community to understand the genetic heterogeneity of ASD (Chen et al., 2018; Darnell et al., 2011; Krishnan et al., 2017).

The *Animal Model* (AM) module of AutDB was created to add depth and precision to the annotation of ASD related models. The genetic models in AM are integrated with the corresponding gene in the human gene module of AutDB, providing human genetic evidence underlying each rodent model. Our database also includes various types of environmentally induced models for autism reported in the scientific literature. Several prenatal factors, including exposure to drugs (Smith and Brown, 2014), role of paternal age (Kojima et al., 2018) and maternal immune factors circulating during gestation (Slawinski et al., 2018) are being studied as causative or modulatory inducers of ASD. Additionally, the complex effects of chemical exposure and drugs after birth are also undergoing scrutiny (Ali and Elgoly, 2013; Foley et al., 2015; Kalkbrenner et al., 2015; Laugeray et al., 2014; Li et al., 2018; Wagner et al., 2006). In contrast to Mouse Genome Informatics (MGI), the paramount resource for mouse genetics for over 35 years, AutDB is a specialized resource that includes diverse types of ASD-related animal models, evidence or hypothesis-based, including inbred strains showing face-validity to ASD, annotated using a shared and standardized framework. Another distinctive and unique feature of the AutDB resource is the inclusion of rescue models, in which drugs and procedural, genetic or dietary manipulations are used in rodent ASD models in an attempt to rescue ASD-relevant phenotypes. Together, AutDB represents a comprehensive resource including genetic and nongenetic animal models relevant in ASD biology.

Using data curated in the AM module over the past 8 years we demonstrate characteristic patterns in analyses undertaken to study genetic and environmentally induced rodent models of ASD. Analysis of trends based on rodent model findings shows that the most frequently assessed phenotypes are related to core features of human ASD such as social interactions, ultrasonic vocalization and repetitive behavior. Additionally, neuroanatomical features like changes in dendritic architecture, observed in post-mortem human studies of ASD brains (Martinez-Cerdeno, 2017), are also frequently examined in rodent models along with electrophysiology conducted on acute brain slices. With a view to facilitating translational research, we highlight pharmaceutical drugs administered to several ASD models. Finally, as a case study, we present a comprehensive analysis of phenotypes studied in rodent models of Shank3, one of the leading genetic risk factors of ASD.

## Results

### 1. Representation of animal models in AutDB

AutDB is an open-access portal designed to provide a comprehensive view of risk factors associated with ASD. Adopting a systems biology approach, this resource integrates diverse functional information of ASD risk-factors while conserving their biological relationships. The AM module develops on the *Human Gene* and *CNV* modules of AutDB by including detailed phenotypic information of animal models. All genetic models are connected to the corresponding factor curated in AutDB for their relevance to ASD (Figure 1). The AM module catalogues induced models based on environmental risk factors and inbred strains with ASD-consistent phenotypes. Information regarding animal models is extracted from published, peer-reviewed primary reports and parsed to provide a detailed view of the constructs and corresponding phenotypic data. The annotation of animal models is guided by a metadata repository of phenotypic terms (phenoterms) and experimental paradigms known as Phenobase (Kumar et al., 2011). Phenoterms are systematically classified into 16 broad categories that align with human ASD phenotypic features (Figure 1b, Supplementary Table 1). Finally, AutDB integrates data from both mouse and rat models using a shared annotation framework for robust analysis.

**Figure 1.**
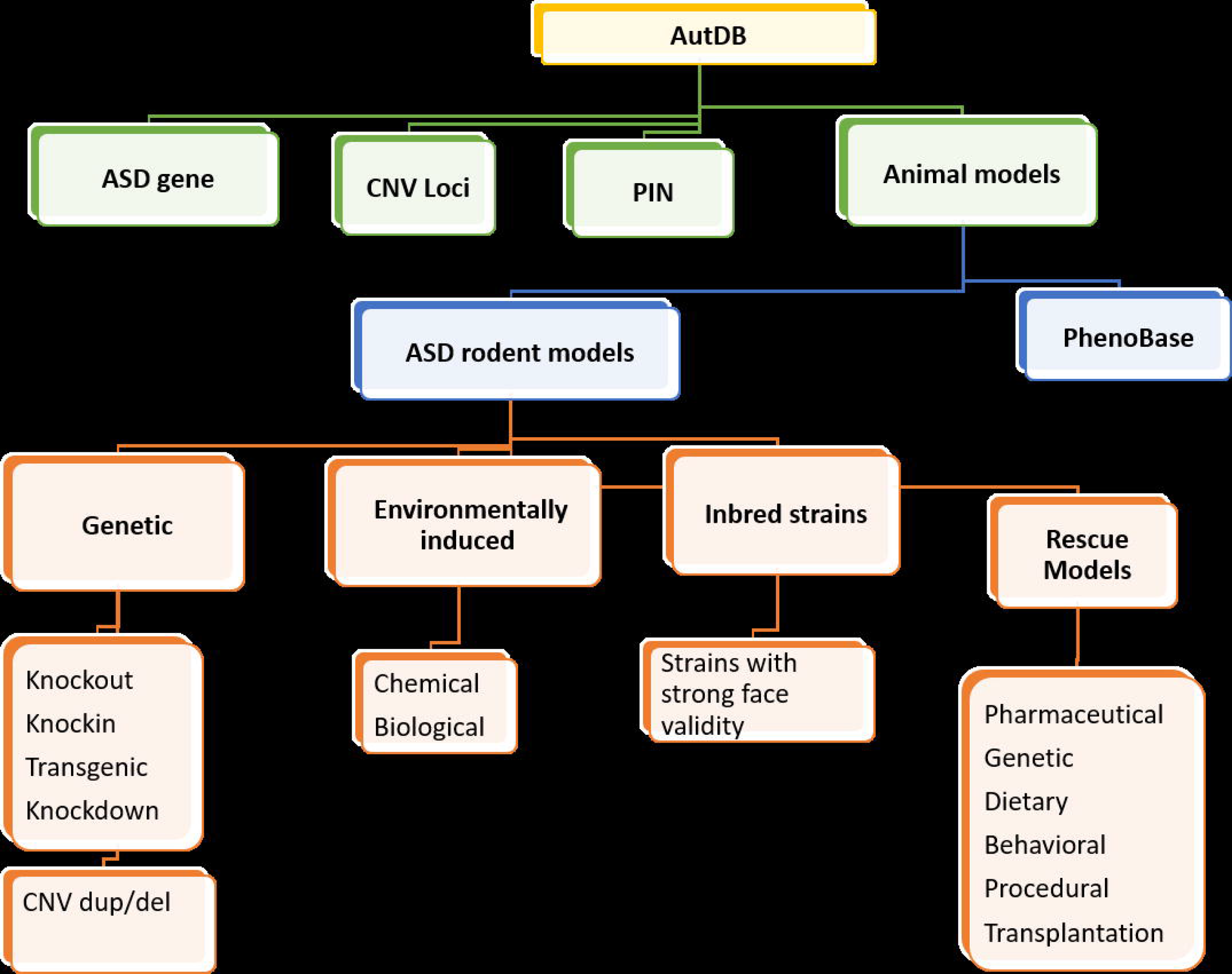
Curation of animal models in AutDB. A. AutDB features a modular framework that aims at collating the multifactorial risk architecture associated with ASD: 1) Human Gene module curates all known human genes linked to ASD together with detailed description of variants associated with the disorder; 2) Copy Number Variant (CNV) module catalogs deletions and duplications of chromosomal loci implicated in ASD; 3) Protein Interactions (PIN) module builds networks of interacting proteins implicated in the etiology of ASD; 4) Animal Model (AM) module collects behavioral, anatomical, and physiological data corresponding to genetic and induced models of ASD. A multilevel data-integration strategy is used to connect animal models to the corresponding entries in the Human Gene and CNV modules, respectively. Inclusion of models originating from relevant non-genetic risk factors and potential quantitative trait loci expand the repertoire of ASD models. Uniquely, AM includes rescue lines based on ASD models which have been treated with an agent to alleviate ASD-related symptoms. B. Frequency of phenotype terminology in phenobase. The number of phenoterms in the Phenobase by category (phenocategory). The categories are qualified as ‘core’, ‘auxiliary’, depending on the extent of relation (accepted by the animal model research community) of the phenotypes (phenoterms) to core or auxiliary endophenotypes observed in human ASD, whereas ‘physiological’ and ‘other’ refer to categories that are assessed in rodents frequently but has no definitive parallels to human ASD.

### 2. Overview of data

The ASD rodent models described in this study were based on over 258 genes, 6 CNV regions, 72 inducers, and 9 inbred strains. These are referred to as ‘ASD factors’ henceforth. Initial analysis indicated that the mouse models were predominantly based on genetic factors, while rat models mainly comprised environmental inducers (Supplementary Figure 1). A wide variation in number of publications per ASD factor was also observed (Figure 2a). The most annotated ASD factors (> 20 articles per factor) included the syndromic genes (Mecp2 and Fmr1) and the inbred strain (BTBR T+ Itpr3tf/J (BTBR)) in mouse (Figure 2a) and valproic acid (VPA) in rat (Figure 2b). ASD models based on face validity such as inbred strain BTBR were among the highly annotated factors reflecting an intense focus of the research community. The next tier of annotation included high confidence *ASD* genes (Shank3, Chd8, Pten, Adnp), syndromic genes (Tsc1, Tsc2, Ube3a), recurrent CNVs (16p11.2, 22q11.2) and nongenetic factors (polyinosinic:polycytidylic acid (poly I:C) and VPA in mouse; lipopolysaccharide (LPS) in rat. Increasing evidence for nongenetic factors linked to ASD etiology, such as maternal immune activation (MIA) has propelled the development of such rodent models. Several mouse and rat models based on MIA induction by agents such as poly I:C, a viral mimetic, or LPS, a bacterial mimetic is represented prominently in the rodent dataset. Finally, a number of environmental factors with uncertain links to ASD such as citalopram, thalidomide, terbutaline, kainic acid, stress and maternal isolation also comprise the rat dataset (Figure 2b).

**Figure 2.**
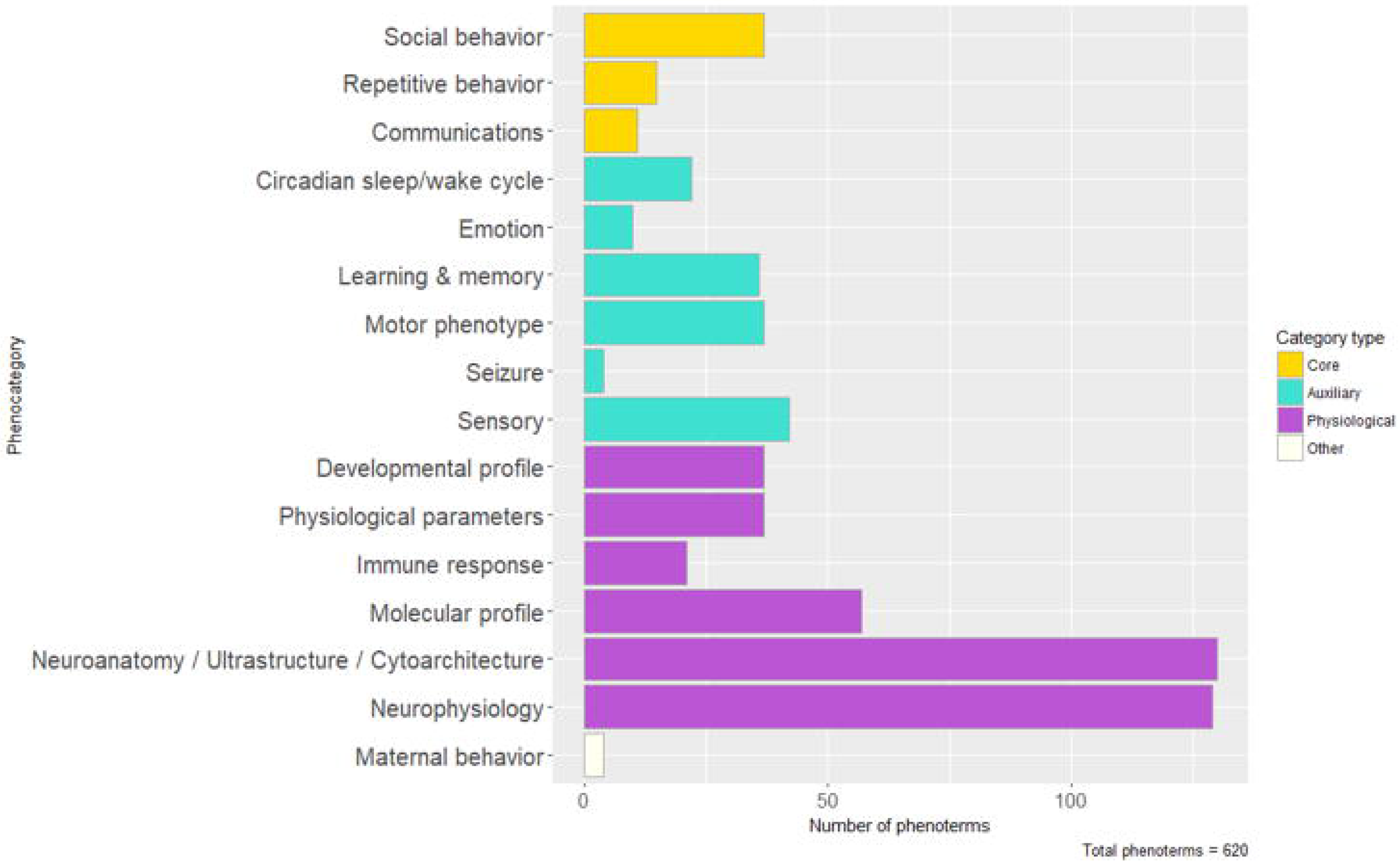
Representation of ASD-associated factors in the rodent datasets of AutDB. A. The number of references and models annotated by the type of ASD factor in the mouse dataset. B. The number of references and models by the type of ASD factor in the rat dataset. The type of the ASD-associated factors is shown by distinct outline colors: gene (red), CNV (pink), induced (blue), and inbred (green). Note that only factors that have four or more references are shown in the mouse (A) and two or more references are shown in rat (B). Abbreviations: VPA, valproic acid; Poly I:C, polyinosinic:polycytidylic acid; APA, advanced paternal age; PPA, propionic acid; LPS, lipopolysaccharide; FRAb, folate receptor antibody; FAST, seizure-prone rat.

### 3. Signature data from ASD rodent models

To explore the data annotated in AutDB, which represents a comprehensive segment of research into ASD etiology, we analyzed the patterns in observations made in rodent models. In all, the data represents phenotypes of rodent models based on 345 ASD factors taken from 787 references. The frequency of phenoterm use across models is a measure of the validity of the phenoterm as a node of comparison. We determined the top 30 most frequently annotated phenoterms in AutDB are distributed across core, auxiliary and physiological categories (Figure 3a). Specifically, we find observed changes in ‘General locomotor activity’, ‘Anxiety’ and ‘Social interaction’ as the three most frequent phenoterms annotated in AutDB.

**Figure 3.**
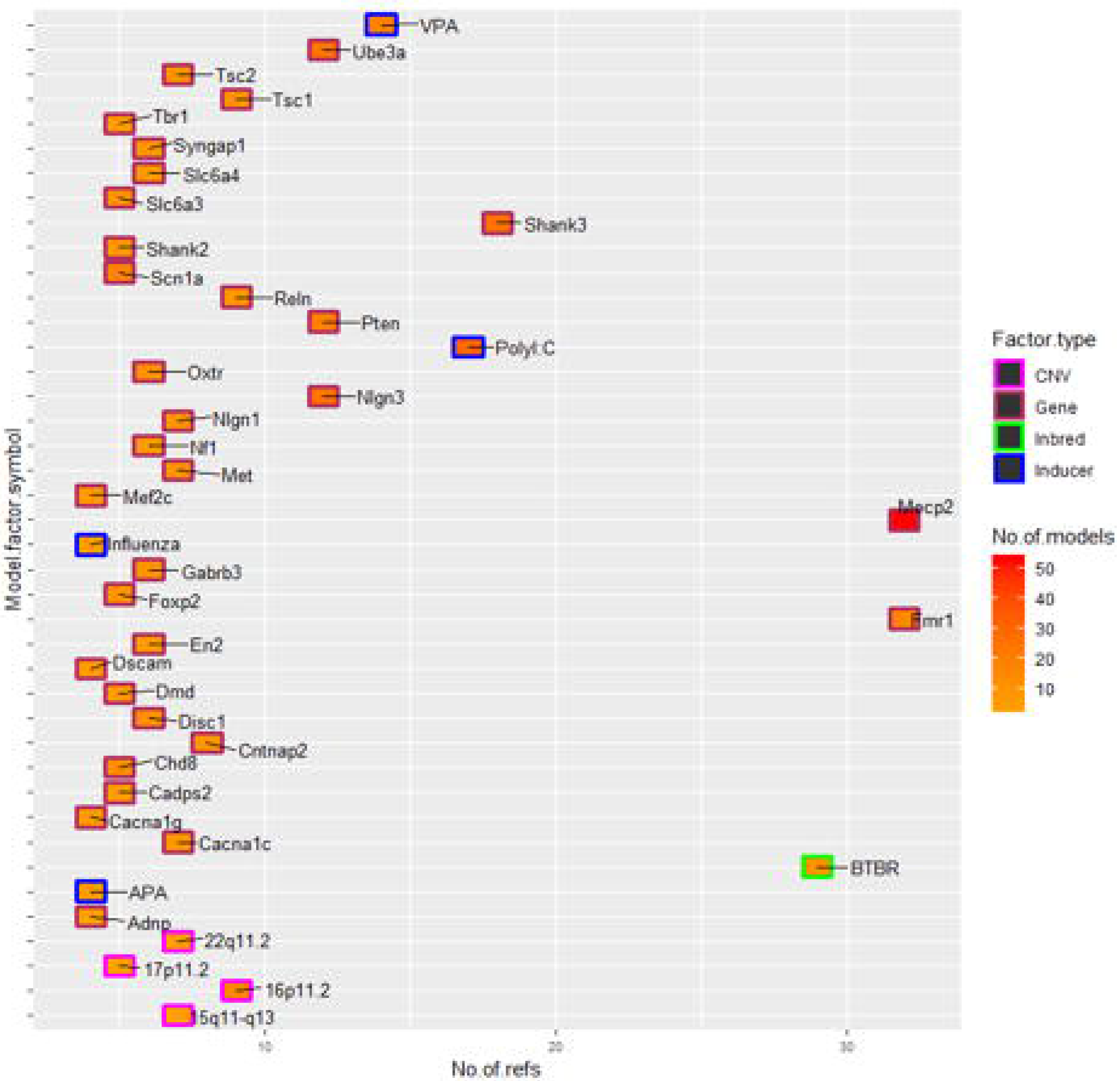
Signature data in AutDB. A. Characteristics of the top 30 phenotypic terms in the rodent dataset. Number of instances of the most frequently annotated phenotypic terms are organized in their respective categories. The categories are color-coded indicating: core phenotypes (ochre), associated phenotypes (green) or physiological observations (purple). B. Changes observed in the most frequently annotated phenotypic terms in rodent datasets. The percentage of total instances of different valences for phenoterms are color-coded: abnormal (purple), decreased (blue), increased(red), and no change (green). Note that the phenocategory ‘Molecular Profile’ is not included in this figure.

In our annotation, phenoterms are paired with a qualitative value term or “phenovalue” to indicate the direction of change compared to control animals. Phenovalues for ASD models can be ‘Increased’, ‘Decreased’, ‘Abnormal’ or ‘No Change’. This is a key feature for building the phenotypic profile of ASD models reported in hundreds of scientific reports. We mapped the percentage distribution of phenovalues for the 30 most frequently used phenoterms (Figure 3b). Phenoterms in core categories showed higher incidence of ASD-consistent measures of phenovalue: ‘Ultrasonic vocalization’ (Decreased = 45%; Increased = 15%), ‘Social memory’ (Decreased = 66%), ‘Social interaction’ (Decreased = 54%), ‘Social approach’ (Decreased = 59%), ‘Self-grooming’ (Increased = 54%) and ‘Repetitive digging’ (Decreased = 17%; Increased = 42%). However, in 14 out of the 30 phenoterms, ‘No Change’ accounted for more than 50% of the annotation for that phenoterm. These mostly represent standard control measures conducted in disease models to assess the validity of more complex behavioral tasks. For example, animals may be tested in ‘Startle response’ prior to testing ‘Cued or contextual fear conditioning’. Some of the most frequently used phenoterms from auxiliary categories are also routinely tested in various disease models of neurodevelopmental disorders, like ‘Spatial learning’. The phenoterms from physiological categories are frequently tested in rodent ASD models with variable outcomes. Interestingly, the phenoterms ‘Synaptic plasticity’ and ‘Synaptic transmission’ are reported as ‘Decreased’ 39% and 42% of the time, respectively, reflecting a heterogeneous contribution of ASD factors towards synaptic function.

### 4. Rescue models

In AutDB, rescue models originate from established ASD models (genetic, induced or inbred) undergoing a treatment protocol with the aim of alleviating one or more ASD related symptoms. A rescue paradigm is defined based on a unique combination of rescue agent, dosage and timing of treatment. Rescue agents are further categorized according to the type of intervention (Supplementary Table 2). In some cases, rescue models provide an understanding of mechanisms underlying ASD related phenotypes whereas in other studies rescue models employs pharmaceutical agents (e.g., FDA approved memantine and rapamycin) to establish or validate their therapeutic use ASD patients.

As noted before, genetic models comprise the most frequently annotated entities in AutDB (Figure 2). Additionally, the majority of rescue paradigms tested on genetic models of ASD are based on pharmaceutical interventions (Figure 4a). Therefore, we focused on this dataset comprising a total of 123 pharmaceutical agents that were identified in various rescue paradigms in AutDB. We further identified 24 drugs used in more than one paradigm indicating the prevalent credence in their putative role in ASD therapy (Figure 4b). Notably, in the list of 32 genes obtained using this prioritization, 10 are key ASD genes with multiple lines of human genetic evidence (SFARI categories 1-2), three are well-known syndromic genes, 8 are genes with suggestive evidence (category 3), and 11 are lower scoring or functional genes where the link to ASD is not fully established from human genetic evidence (Supplementary Table 3). The correlation between the pharmaceutical rescue agents and the targeted ASD rodent models are shown in figure 4b. Given the importance of altered E:I ratio in ASD (Rubenstein and Merzenich, 2003), several drugs based on the common molecular substrate of glutamate receptors (N-methyl D-aspartate (NMDA) or metabotropic (mGluR) were tested on multiple genetic models of ASD (Supplementary Table 4). Interestingly, three different drugs functioning as agonists for gamma aminobutyric acid receptor subunit A (GABA-a) rescued ASD-related symptoms in multiple models arising from important ASD genes (SHANK2, SCN1A, GABRB3, GRIN1, UBE3A, ARHGAP32). Some of the other frequently tested drugs (fluoxetine, memantine, 2-methyl-6-(phenylethynyl)pyridine (MPEP), lithium) are currently prescribed for psychiatric disorders However, a small subset of drugs was only tested on models based on a single gene: JQ1 and FRAX48 in Fmr1 models; NAP in Adnp models; and p-cofilin in Shank3 models. In these cases, the drugs targeted specific pathways regulated by the ASD gene. For example, JQ1 is an inhibitor of bromo and extra terminal domain (BET) proteins that are regulated by fragile-X mental retardation protein (FMRP).

**Figure 4.**
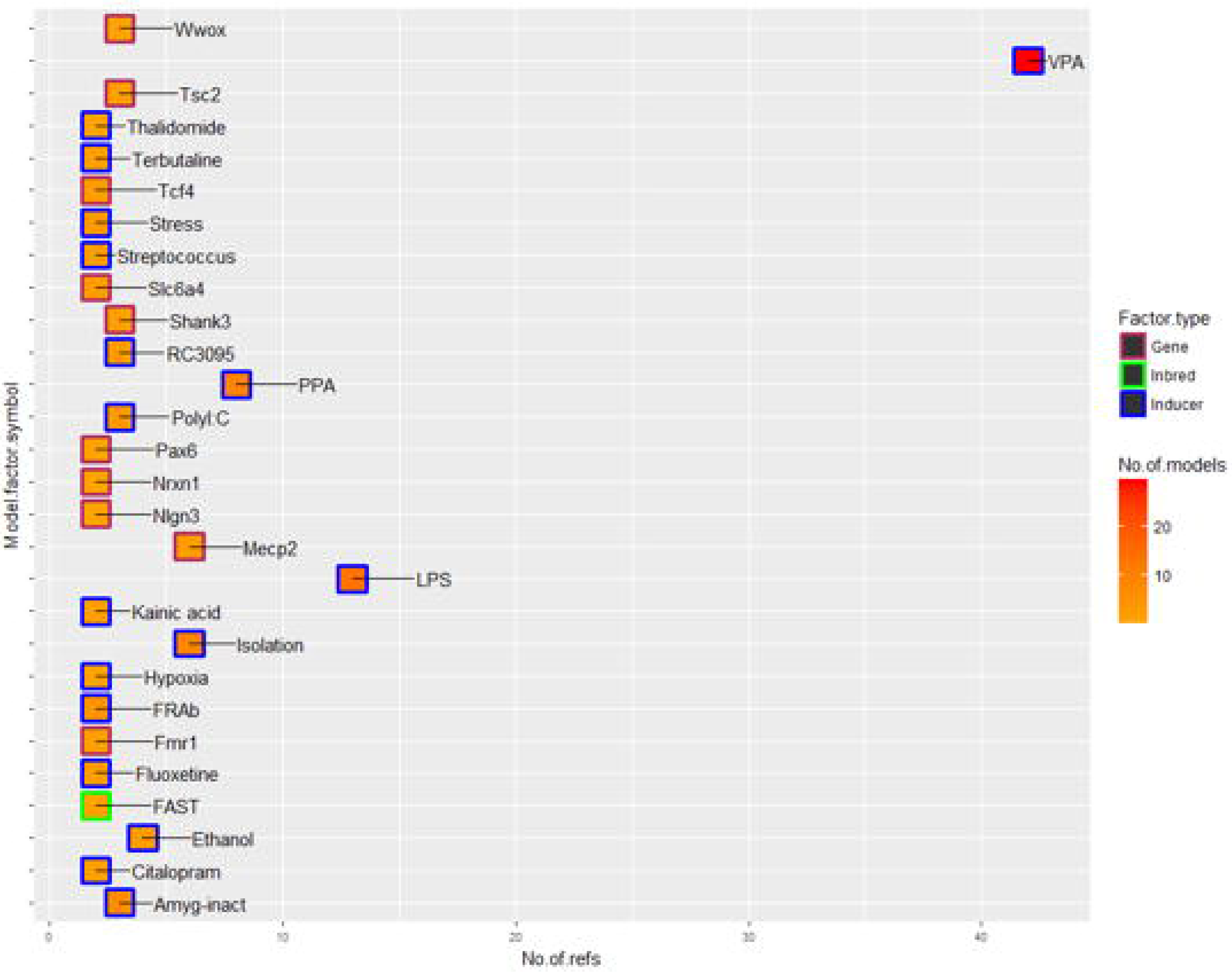
Pharmaceutical agents used in alleviating ASD-related symptoms in rodent models. A. The pie-chart shows the percentage of rescue paradigms based on different types of interventions: pharmaceutical agents, genetic, dietary or procedural tested on genetic rodent models of ASD. B. This scatter plot depicts the pharmaceutical agents tested on rodent ASD models based on the genes shown in the X axis. Y-axis indicates name of the rescue agent. Only agents that were tested in more than one rescue paradigm are shown. ‘n’ refers to the number of treatment paradigms tested on models based on the same gene.

### 5. Case Study: Shank3

While studying the overall phenotypes in a large set of ASD models provided an overview of ASD literature and trends in research, an in-depth analysis of an important ASD-linked genetic factor, Shank3, illustrates the functionality of the intricate annotation in AutDB. Shank3, a multi-domain scaffolding protein with a prominent role in glutamatergic synapses has been implicated in ASD through the identification of rare damaging mutations in multiple studies (high confidence SFARI Gene). Accordingly, several research groups have reported mouse models of Shank3 in an attempt to define its contributory role in ASD. However, the complex structure of the Shank3 gene with multiple exons and experimentally validated alternative splicing from intragenic promoters presented a major challenge in establishing relevant animal models. The regions targeted in mouse models that are annotated in AutDB are shown in Figure 5a. Overall, there are 27 loss-of-function (LOF) models of Shank3 including 15 knockout (KO) and 10 knockin (KI) mouse models together with two KO rat models (Supplementary Table 8 and 9 for details on domains targeted and model constructs). Targeting these protein domains in Shank3 resulted in different sets of isoforms being lost, further adding to the resultant phenotypic complexity (Monteiro and Feng, 2017; Wang et al., 2014).

**Figure 5.**
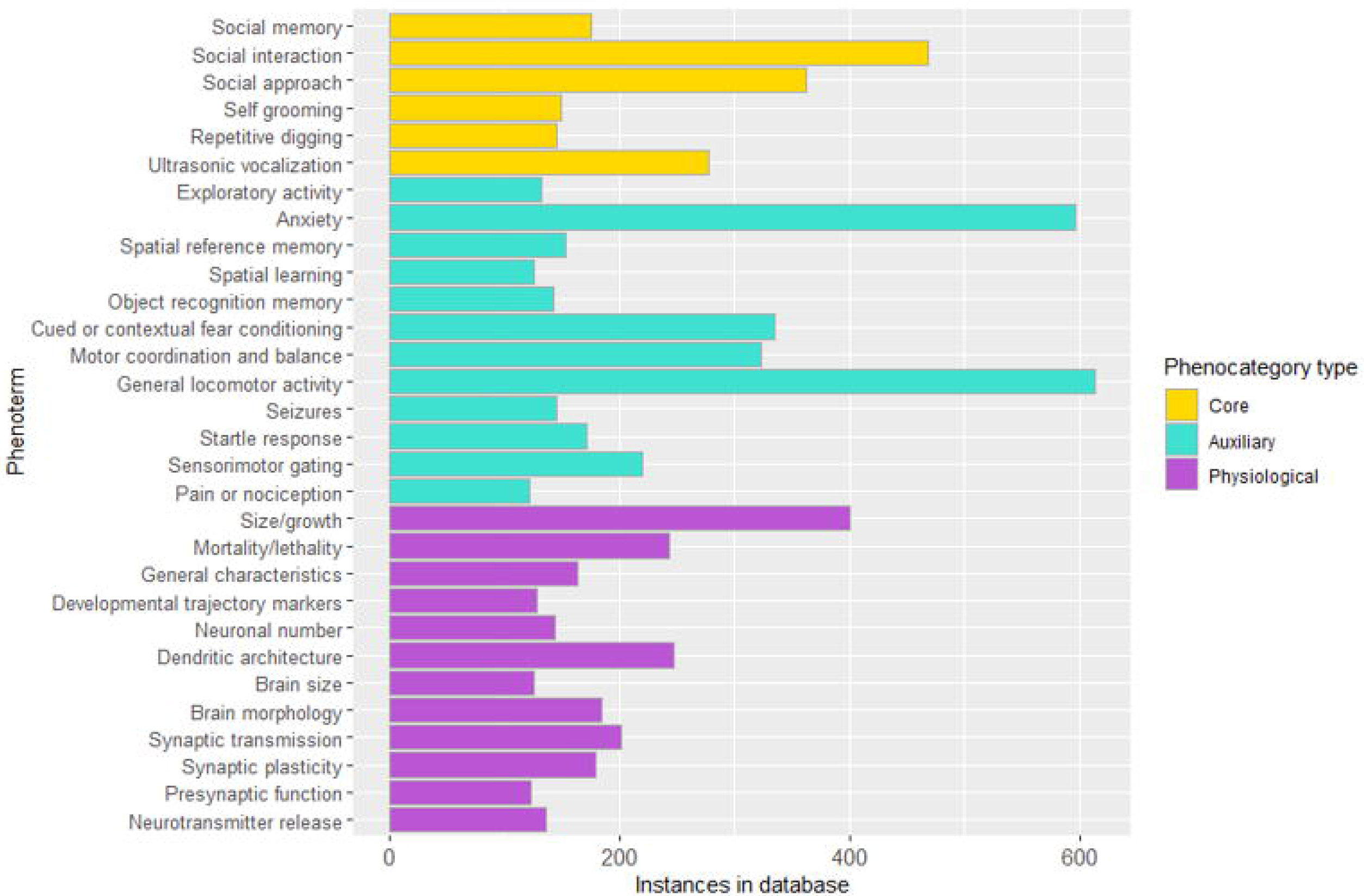
Shank3 model characteristics displayed by domain targeted. A. Schematic representation of the relative positions of exons (blue vertical lines) and protein domains in mouse Shank3, with numbers marking exons containing the respective domains (indicated by brackets on top). The solid horizontal black lines indicate the targeted regions in Shank3 KO mouse model constructs present in AutDB. The arrow indicates the approximate location of KI constructs, black arrows are human mutations replicated in mouse. Relative positions of introns and exons based on Ensembl. This graphic depicts the most frequently assessed phenotypes of Shank3 mouse models present in our database, displayed here separately for all homozygous (HM) KOs genotypes. Each tile color represents the phenovalue and phenotypes have been arranged in a grid by the domain targeted in the KO constructs. C. Detailed view of core phenotypes of Shank3 HT and HM models based on ANK, PDZ and PRO targeted domains. This figure depicts that several core phenotypes are exhibited by HT and HM models and shows the presence of apparently contradictory data in behavioral analyses, which in some instances can be explained due to differences in age or sex of animals tested (see main text).

Unlike many ASD genes essential for survival (Ji et al., 2016), homozygous (HM) loss of Shank3 is compatible with life through developmental stages to adulthood in both mouse and rat. Consequently, a broad range of assays targeting ASD-related core and associated phenotypes were performed in various Shank3 models (Supplementary Figure 2). The most frequently assessed phenotypes in Shank3 mouse models were similar to the overall phenotype reporting trends in the entire set of ASD models in AutDB (Supplementary Table 5). Both ‘Anxiety’ and ‘Self-grooming’ were increased in all tested HM KO mouse models (Figure 5b) indicating a strong recapitulation of human ASD phenotypes in Shank3 KOs. Similar in essence to the allelic heterogeneity in human ASD phenotypes (Leblond et al., 2014) a wide variability of phenotypes in Shank3 KO models was observed based on the targeted domain. Surprisingly, full-length KOs displayed normal social behavior whereas some deficits in social behavior were seen in models targeting ANK, PDZ, and PRO regions. Ultrasonic vocalization (USV), another ‘core’ phenotype postulated to assay for communicative behavior, was also variably affected in the different Shank3 models, with full-length and ANK targeted models displaying impairments in USV calls while models with mutated PDZ domains showed normal USV phenotype. Dendritic spine density in the brain, another aberrant phenotype in ASD patients, was found to be anomalous in all HM Shank3 KO mutants (Figure 5b; AutDB). An expanded view of the top 5 ‘core’ phenoterms indicates that not all phenotypes have been tested in individual constructs. Interestingly, Shank3 heterozygous (HT) mutants manifest some ASD phenotypes indicating that dosage and nature of knocked-out Shank3 isoforms are essential for normal function (supplementary figures 3, 4 and 5). Two human mutations in exon 21 have also been recreated in Shank3 KI models the ASD related InsG and the Schizophrenia (Schz) related R1117X (Speed et al., 2015; Zhou et al., 2016). As shown in Figure 6, compared to exon 21 KO mutants, the KI mutants have similar changes in anxiety and the InsG KI mutants also show similar changes in synaptic transmission to the KOs. Observations from rat models of Shank3, recently added to the AM module indicate that rats and mice display differential changes in behavioral phenotypes (Supplementary Figure 6).

**Figure 6.**
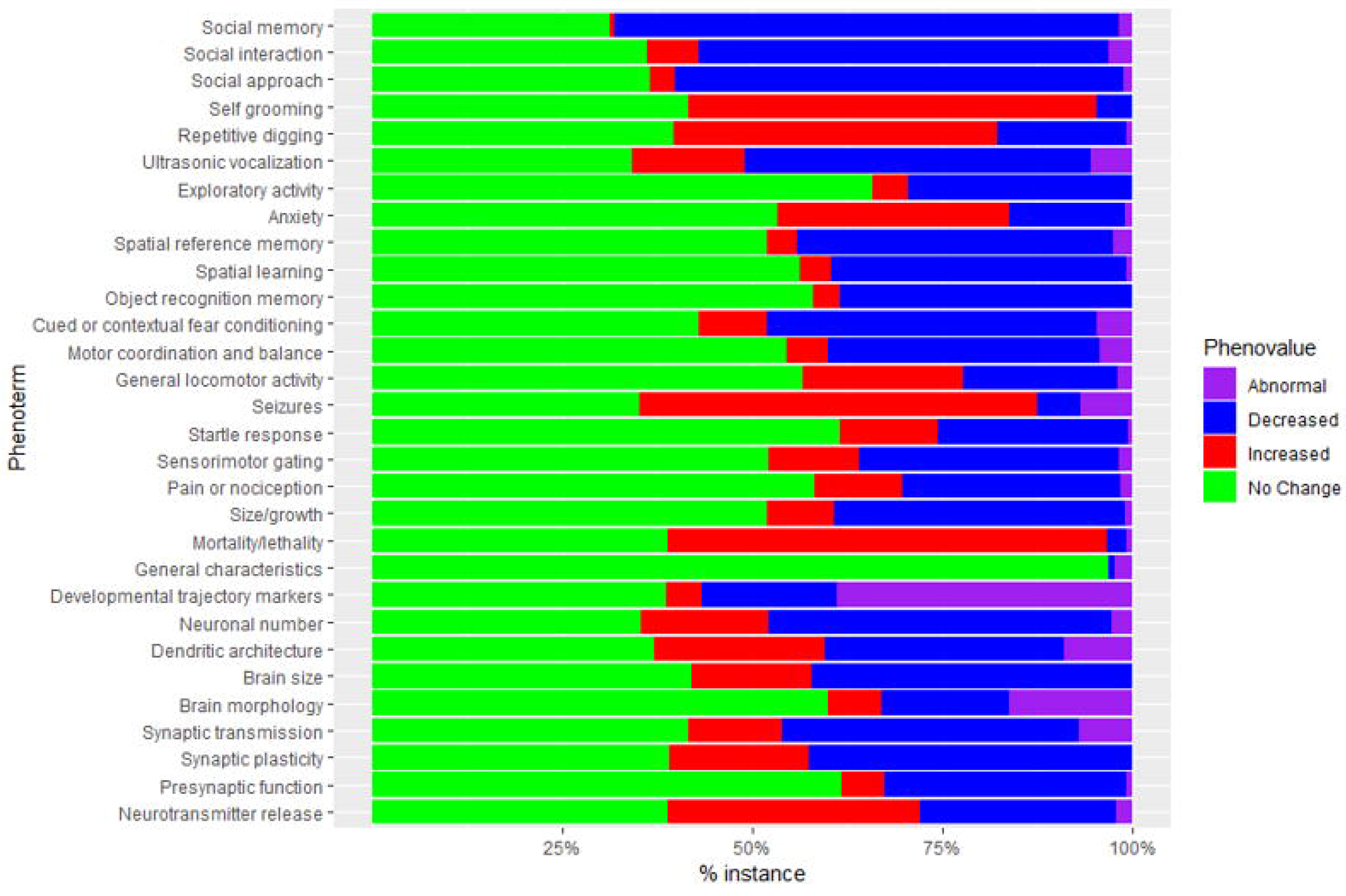
Shank3 PRO domain targeted HM KI and KO models. Phenotypic observations from Shank3 HM KI models by insertion type. ASD(insG) and Schz(R1117X) are human mutations in exon 21 found in people with ASD and Schizophrenia. Social behavior is affected in ASD and Schz related mutations, as well as exon 21 KO that causes the production of fewer Shank3 isoforms. Schz related mutation does not cause an increase in self-grooming, unlike the InsG, including ASD related InsG. Other learning and memory phenotypes, neurophysiological and anatomical features are affected in both ASD and Schz specific KI mutation as well as exon 21 KOs.

While several rescue paradigms have been tested on Shank3 models, they were all based on eight LOF models. A genetic approach based on reinstatement of wildtype Shank3 expression after birth resulted in alleviation of a subset of ASD phenotypes (Supplementary Table 6).

Pharmaceutical agents like MPEP, TG003, p-cofilin and CDPPB (3-cyano-N-(1,3-diphenyl-1H-pyrazol-5-yl) benzamide), have been used in different Shank3 models (Table 1). High doses of p-cofilin, TG003, and MPEP restored or ameliorated repetitive self-grooming seen in the Shank3 mutants. Interestingly, p-cofilin and TG003 also normalize social behavior with a sustained effect (Supplementary Table 7). Rescue attempts on rat Shank3 models have in turn utilized oxytocin, which restored some deficits in social behavior and reward reinforced choice behavior (Table 2). Furthermore, neuroreceptor activity and synaptic neuroreceptor based transmission were restored in ANK and PRO domain targeted Shank3 mutants treated with p-cofilin and IGF-1. In the rat Shank3 model, synaptic plasticity is found to be completely restored in the medial prefrontal cortex and partially restored in the hippocampus following intracranial injection of oxytocin, indicating an avenue for translational studies.

**Table 1.** Restored or ameliorated phenotypes in Shank3 mouse models displaying the targeted domain(s), the experimental paradigms and drug name.

| Domain | Phenoterm | Experimental paradigm | Effect | Agent | Citation |
| --- | --- | --- | --- | --- | --- |
| All | Reward reinforced choice behavior | Operant conditioning paradigm | Ameliorated | CDPPB | (Wang et al., 2016) |
|  | Protein localization: synapse | Western blot: striatum | Ameliorated |  |  |
|  | Synaptic plasticity: striatal LTD | Whole-cell patch clamp | Restored |  |  |
|  | General locomotor activity | Open field test | Restored | MPEP |  |
|  | Self grooming: perseveration | Grooming behavior assessments | Ameliorated |  |  |
| ANK | Motor coordination and balance | Accelerating rotarod test | Restored | IGF-1(high dose) | (Bozdagi et al., 2013) |
|  | Neuroreceptor activity | Whole-cell patch clamp | Restored |  |  |
|  | Synaptic plasticity: hippocampal LTP |  | Restored |  |  |
|  | Neuroreceptor activity |  | Restored | peptide derivative of IGF-1 |  |
|  | Synaptic plasticity: hippocampal LTP |  | Restored |  |  |
| PRO | Synaptic transmission: excitatory | Whole-cell patch clamp | Ameliorated | constitutively active Rac1 | (Duffney et al., 2015) |
|  | Social approach | Three-chamber social approach test | Ameliorated |  |  |
|  | Cytoskeletal organization | Western blot: actin and F-actin levels | Restored | p-cofilin peptide (high dose) |  |
|  | Self grooming: perseveration | Grooming behavior assessments | Restored |  |  |
|  | Synaptic neuroreceptor ratio (NMDAR/AMPA) dependent transmission | Whole-cell patch clamp | Restored |  |  |
|  | Synaptic transmission: excitatory |  | Restored |  |  |
|  | Social approach | Three-chamber social approach test | Restored |  |  |
|  | Protein phosphorylation | Western blot | Restored | TG003, CLK2 inhibitor | (Bidinosti et al., 2016) |
|  | Self grooming: perseveration | Grooming behavior assessments | Ameliorated |  |  |
|  | Social approach | Three-chamber social approach test | Restored |  |  |

**Table 2.** Restored or ameliorated phenotypes in Shank3 rat models displaying the targeted domain, the experimental paradigms and drug name.

| Domain | Phenoterm | Experimental Paradigm | Effect | Agent | Citation |
| --- | --- | --- | --- | --- | --- |
| ANK | Reward reinforced choice behavior | Operant conditioning paradigm | Ameliorated | Oxytocin | Harony-Nicolas et al., 2017 |
|  | Social memory: long-term social memory | Reciprocal social interaction test | Ameliorated |  |  |
|  | Synaptic plasticity: hippocampal LTP | Field potential recordings | Ameliorated |  |  |
|  | Synaptic plasticity: mPFC LTP | In vivo local field potential (LFP) recordings | Restored |  |  |

## Discussion

AutDB is a platform designed to be a specific resource where the AM module focuses on the in-depth annotation of genetic and non-genetic ASD models, using multiple layers of standardized vocabulary encapsulated in the phenobase. Therefore, our database encompasses models based on high confidence ASD genes (e.g., Chd8, Shank3), environmental inducers (e.g., VPA, MIA via exposure to viruses or viral mimetics like polyI:C) and inbred strains (e.g., BTBR). Rat models of ASD were recently added to AutDB to exploit interspecies conserved biology in investigating genetic and environmental ASD risk factors, also believed to be the best approach in the success of clinical trials. Additionally, rats are used in more studies assessing the effects of inducers, thereby increasing our repertoire of inducers tested in rodent ASD models. As the exact genetic signature of ASDs still remains to be determined and is a field of intense ongoing research, we believe that including putative and established models that represent characteristics of autism will lead to a more comprehensive understanding of this complex disorder.

At the time of data freeze for this article (March 2018), AutDB included rodent models based on 258 genes linked to ASD, linked to our *Human Gene* module providing details of all rare and common variants in these genes identified in affected individuals. In contrast, mouse models based on only 57 genes have been linked to ASD in MGI (August 2018). In addition to providing ASD-specific genetic relevance to the animal models, a number of features distinguish our annotation model from MGI. First, our integrative approach includes genetic and nongenetic models of ASD within a single platform. Second, a standardized phenotypic repository (Phenobase) structured according to the diagnostic symptoms of ASD guides annotation of all animal models in AutDB. Third, we make dedicated provisions for all the confounding details that can give rise to contradictory data and uphold robust findings. AutDB provides distinct information on the experimental paradigms used to assess a phenotype and relevant experimental details specific to studies, information that is not available in MGI. One of the primary confounders is the experimental paradigm used to assess a phenotype, e.g., anxiety can be assessed in as many as five to six direct testing protocols, including open-field test, light-dark exploration, elevated plus maze and novelty-induced hypophagia (Bailey, 2009), while being reported as auxiliary observations from several other protocols like Morris water-maze, social behavior testing etc. (Larrieu et al., 2017; Schulz et al., 2007). As noted in results, anxiety is one of the most frequently assessed phenotypes in ASD models as well.

In our unique rescue model dataset, we highlight test outcomes from experimental drugs, FDA approved drugs, behavioral interventions, and transplantations of remedial cells or microbes, on rodent models of ASD. We also curate genetic manipulations that have been used to rescue ASD related phenotypes. Genetic, behavioral or transplantation-based rescue paradigms play an important role in understanding the contribution of different factors for the development of phenotypes in ASD, even if not all of them can be translated into human interventions. We aim to curate new technologies as their use in ASD research is increasing, however, caveats in their application emerge retrospectively, e.g., clozapine-N-oxide (CNO) used to activate designer receptors exclusively activated by designer drugs (DREADDs) is reverse metabolized clozapine (interestingly a drug tested in ASD models) in rodents and is not pharmacologically inert (Manvich et al., 2018). Therefore, it is of great interest to the scientific community to combine the time-tested standard tasks and sophisticated frontier technology to unveil interesting relationships between different types of behavior.

The rescue model dataset of AutDB offers a comprehensive resource for translational studies. For example, genetic rescues on Shank3 models of ASD (see Supplementary Table 6) indicate the crucial role of normal Shank3 expression through adulthood, indicating the importance of these Shank3 mutant lines in testing remedial approaches for Shank3 mutations during postnatal development and adulthood (Mei et al., 2016). A faster and cost-effective route for new clinical therapy in ASD is the use of existing pharmaceutical agents, or “repurposing” of drugs already approved by FDA for other conditions. As seen in Figure 4 several known drugs have been tested in genetic models, including rapamycin, fluoxetine, clozapine and risperidone. Rescue treatment paradigms and effects, annotated discretely for rescue models in AutDB with a new set of controlled vocabulary distinct from parent ASD models, are as important as the drugs used for rescue as dosage and length of treatment can significantly change the outcome in rodents and people. This is evident in the Shank3 rescue models where a drug, p-cofilin, tailored to act on a deficit specifically manifested in Shank3 mutants: dendritic spine formation displayed differences in outcome in low (0.15 picomol/g) versus high (15picomol/g) doses. Some of the drugs tested in Shank3 models have also been tested in other mouse models of ASD: CDPPB has been used in rescue paradigms to treat Shank2 mutants leading to normalization of social interaction (Won et al., 2012).

Translation from preclinical to clinical studies requires rigorous testing of a large number of paradigms, with changes in dosage and route of administration requiring. While not reported as often as favorable effects, presence of adverse effects of drug treatments are noted in AutDB and could play an important role towards clinical drug trials. We believe refractory phenotypes should be reported whenever testing is conducted, so that the community can benefit from the knowledge and reduce unnecessary waste of resources. From the Shank3 dataset itself, it is surprising that the assessments for perseverative self-grooming and social approach or interaction were not reported after several treatments (Table 1). Overcoming this ‘positive data bias’ is the basis for our comprehensive annotation of the ‘No Change’ phenotype which indicates the absence of ASD related phenotype or no statistical difference from control measures. A single model for ASD will likely not emerge from rodent studies for this genetically and clinically heterogeneous disorder. Similarly, it is unlikely that there will be a single drug prescribed for ASD therapy. As presented here, a number of the most frequent ASD related phenotypes, like social behavior and anxiety, are assessed in Shank3 models. Additionally, based on the scaffolding function of the Shank3 at the synapse, other phenotypes like synaptic plasticity, glutamate neuroreceptor levels and more detailed neurophysiology are also reported. This is where the advantage of rodent models lie: in deciphering the constellation of phenotypes that group together. For example, a recent study using optogenetics indicates that spatial learning can have direct effects on social behavior (Murugan et al., 2017) and it is likely that future research from different fields will enhance the existing knowledge on ASD biology with new insights. Our database is primed to curate new types of paradigms and knit together robust, cumulative observations with new discoveries. Distinctive patterns are yet to be found in ASD research, however it is our aim to provide the platform where in-depth review of ASD scientific literature and standardized annotation reduces confounding factors reporting of all observations in an unbiased manner, to facilitate their detection.

## Materials and Methods

### Data curation and annotation

AutDB consists of manually curated and annotated data from published, peer-reviewed scientific literature on the basis of relevance to ASD. For curation in the Animal Model module of AutDB, articles are selected for annotation based on a preliminary assessment of the validity of the rodent model and the detailed phenotypic characterization of the model. Our database is updated quarterly with annotations from the latest published literature. Details of the annotation process are documented in a wiki web resource (http://174.79.186.155:18000/AM_wiki/index.php/Rodent_annotation_guideline).

The validity of an animal model is based on its relevance to ASD. The articles must describe animal models that are based on evidence from association studies in humans, or, alternatively, models that display strong face validity for ASD-consistent endophenotypes. An animal model where an ASD-associated factor is manipulated to assess resulting phenotypes is an evidence-based model, while a model showing ASD-consistent endophenotypes for a factor with no association with ASD is a hypothesis-based model. We attempt to build a comprehensive dataset for rodent models based on high confidence ASD genes and prioritize reports containing detailed phenotypic data on those. Additionally, rescue models are annotated based on rescue paradigms where an agent or intervention is used to alleviate a phenotype in an ASD animal model. We categorize these rescue paradigms based on the type of agent: transplantation-based, procedural, behavioral, genetic or pharmaceutical. We have developed annotation methods to clearly represent the treatment effects on ASD phenotypes paralleled in rodent models.

The data freeze date for all data shown in this article is March 31, 2018.

### Phenobase: dynamic hierarchical phenotypic metadata

The individual phenotypes observed in ASD models are annotated using controlled vocabulary (‘phenoterms’) and organized into 16 broad categories in a resource termed as the Phenobase (Supplemental Table 1). These categories are grouped as ‘core’ where the comprising phenoterms closely parallel ASD core phenotypes, ‘auxiliary’ when the parallel human ASD phenotypes are not core diagnostic features of ASD, or ‘physiological’ for most other associated phenotypes that are routinely assessed to determine biological underpinnings of ASD.

Phenoterms, which are arranged in a hierarchical manner, are based on endophenotypes observed in rodent models. As many complex endophenotypes of rodent animals are being reported, we have added phenoterms that capture specificity while still rooted in a broader term, such that broad terms at the top of the hierarchy are separated by colons from specific terms. For example, the term ‘Morphology of the basal ganglia: Striatum: Caudoputamen’ specifies the morphological changes to the caudoputamen, a part of the striatum, which in turn is a part of the basal ganglia. Our phenoterms are intended to capture complex as well as simple endophenotypes that are physiological, robust or quantitative and conserved between species. It should be noted here that the phenoterms have been developed in the context of curated ASD literature, therefore they reflect the phenotypes assessed in ASD rodent models and not the full complexity and scope of a category per se.

In addition to phenoterms, the Phenobase contains a standardized list of experimental paradigms. This list is a discrete part of the database that is used to represent the tests used to study phenotypes in different animal models. The standardization of experimental paradigms adapts a uniform nomenclature that circumvents the variety of synonyms used in literature for similar experimental set-ups. The Phenobase catalogues over 450 experimental paradigms that map to the whole set of 620 phenoterms. The combined use of phenoterm and experimental paradigm provides a more comprehensive picture of observations made by authors, which allows for better comparisons of model phenotypes between different ASD models as well as between researchers.

### Data analysis and visualization

Most data analysis and frequency measurements were conducted in R using packages: ‘tidyverse’, ‘dplyr’ and ‘forcats’ (Wickham, 2014). Graphics were developed with ggplot2 or in excel (Wickham, 2016).

## Acknowledgments

Authors would like to thank the entire MindSpec team for fruitful discussions during the development of the manuscript. We thank Simons foundation for their generous support.

## Availability of data and materials

All data described in this manuscript will be available in AutDB (http://autism.mindspec.org/autdb/Welcome.do) and SFARI Gene (https://gene.sfari.org/autdb/Welcome.do).

## Author contributions

ID: methodology, software, validation, formal analysis, data curation, writing-original draft preparation and visualization
ME: data curation, writing-review and editing
AS: data curation, writing-review and editing
SB: conceptualization, writing-original draft preparation, supervision and funding acquisition

## Tables and Figures

**Table 1. Phenotypes in Shank3 rodent models restored or ameliorated by pharmaceutical agents.**

## Supplementary Figures

**Supplementary Figure 1.**
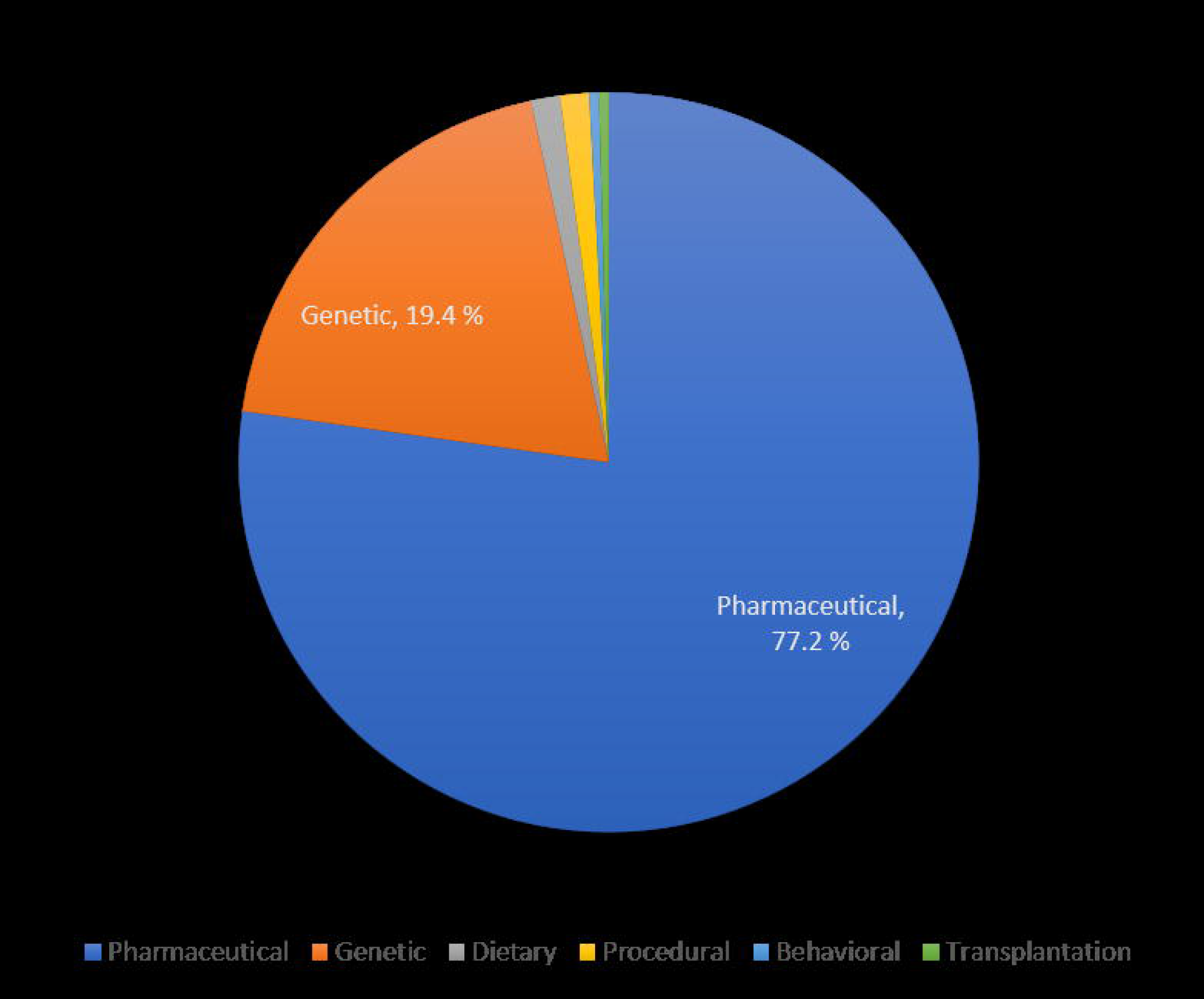
Distribution of ASD factor in rodent data. A. The stacked plot shows that the genes comprise the largest subtype of ASD factors in AutDB with 258 total genes present in the dataset, with most of the models developed in mice. On the other hand, inducers (72) are over represented by rat models and there are about equal numbers of inbred strains that show face validity to ASD in both species. The same ASD related gene or inducer has been modeled in mice and rat infrequently, with only 19/258 genes and 8/72 inducers modeled in both.

**Supplementary Figure 2.**
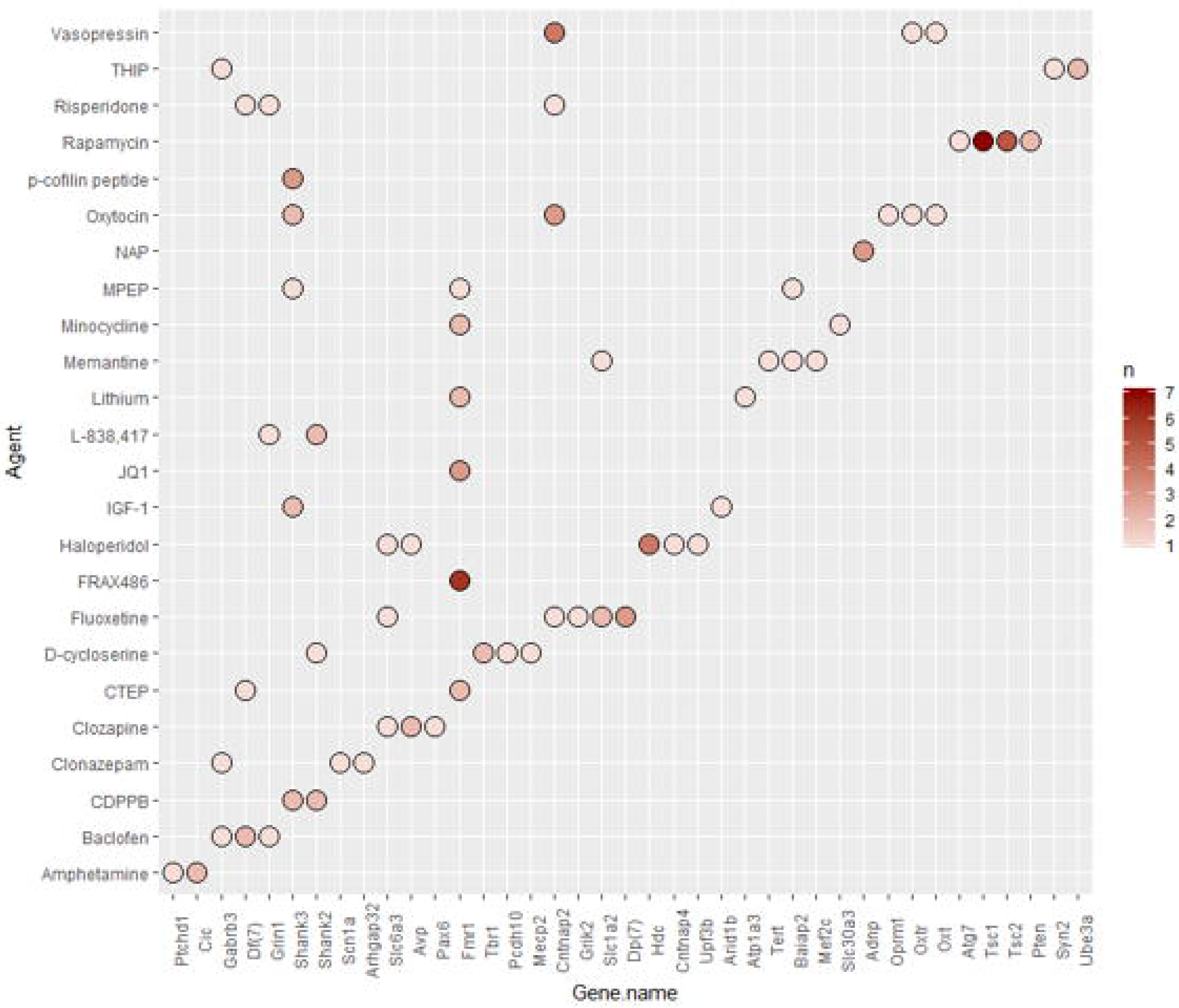
Shank3 model phenotypic data displayed by genotype. An overall representation of Shank3 mouse data separated only by genotype. This figure illustrates that both HM and HT Shank3 KO and KI models have been tested for many phenotypes using different constructs designs (Supplementary Table 8)

**Supplementary Figure 3.**
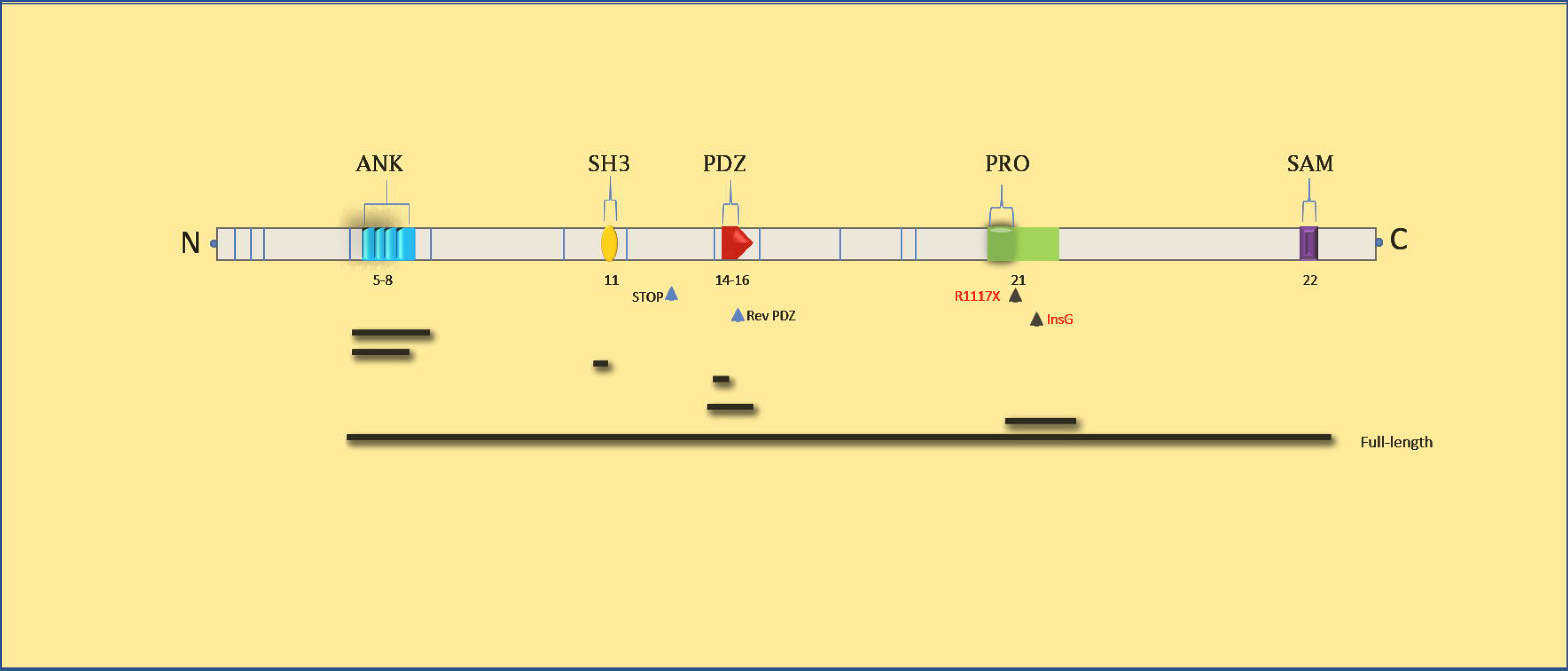
accompanying Fig. 5B. Shank3 heterozygous KO model data depicted by protein domain targeted and genotype.

**Supplementary Figure 4.**
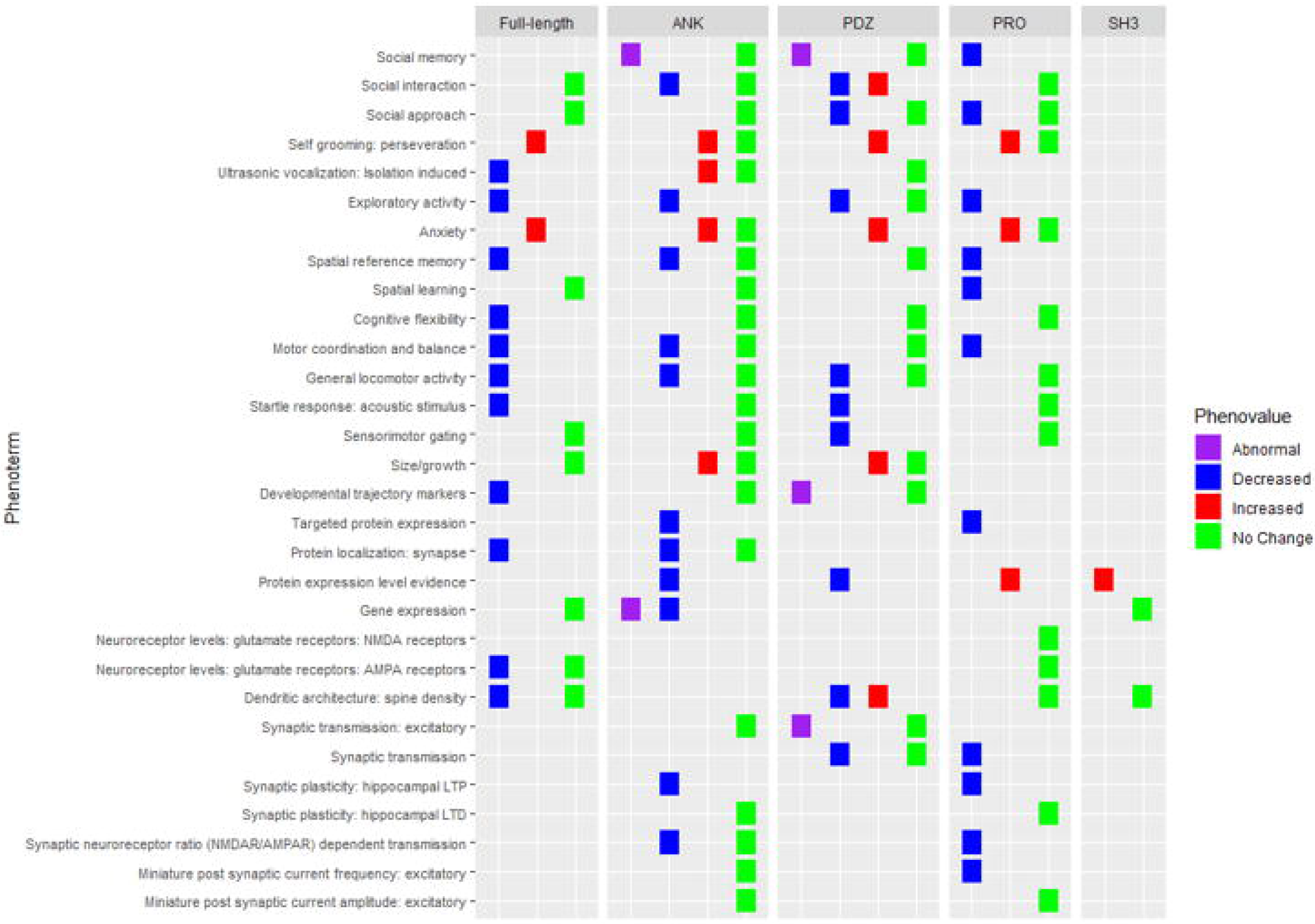
accompanying Fig. 6. Shank3 PRO domain targeted heterozygous KI and KO model data.

**Supplementary Figure 5.**
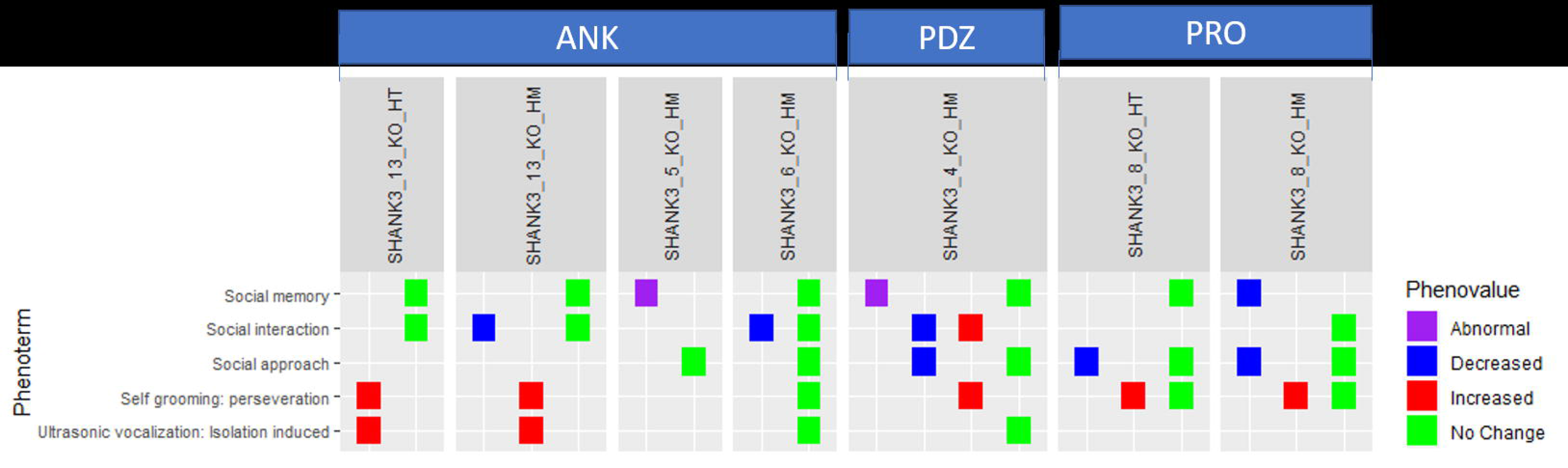
Phenotypes of Shank3 KI models. **A**) The HM KI models display several core phenotypes including impaired social behavior and increased self-grooming. Depending on the mutation, there is heterogeneity in the manifestation of other behavioral phenotypes like anxiety and spatial learning. The KI human mutations are discussed in main text. B) Shank3 HT KI mutant mice still manifest core phenotypes whereas other tested behavior is more like wild type mice, like normal anxiety, sensorimotor gating and spatial learning.

**Supplementary Figure 6.**
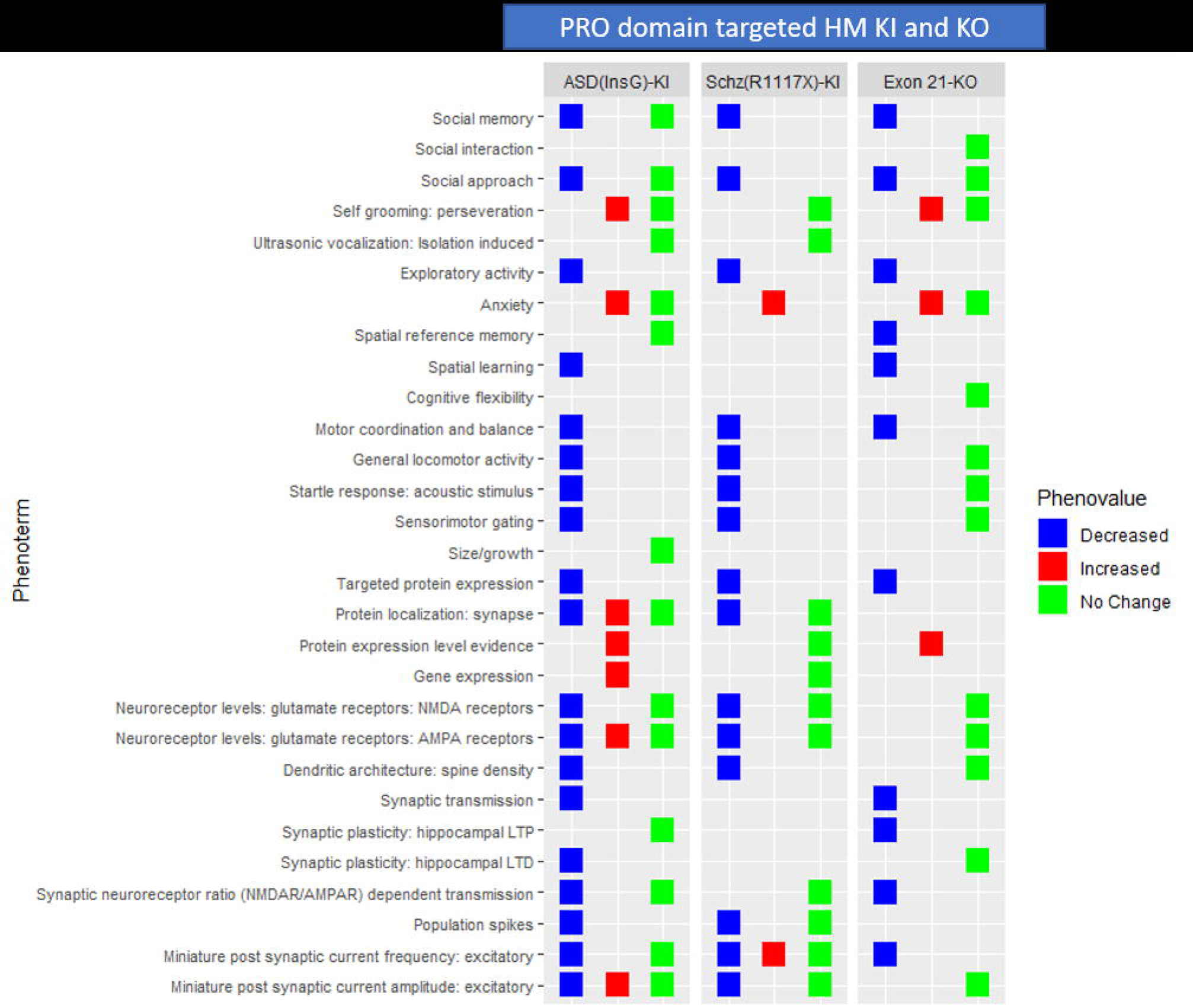
Rat Shank3 phenotypic data. The HT and HM rat models depicted here were developed by targeting the Ank domain. Rat models of Shank3 display some deficits in social behavior (long term memory) but do not share several of the phenotypes displayed by mouse Shank3 models, like no impairments in ultrasonic vocalization or change in anxiety levels are observed in rats.

### List of Supplementary Tables

1. Phenocategory definitions
2. Rescue model type definitions
3. SFARI scores of genes in figure 4b
4. Target or mechanism of 24 drugs tested in 2 or more paradigms on rodent genetic models
5. Shank3 top 31 most frequent phenotypes including targeted protein expression
6. Shank3 rescue data genetic reinstatement
7. Shank3 rescue data all outcomes of drugs
8. Shank3 domains with model IDs
9. Shank3 model construct definitions (all KO and KI) (excel sheet)

## REFERENCES

Ali, E.H., and Elgoly, A.H. (2013). Combined prenatal and postnatal butyl paraben exposure produces autism-like symptoms in offspring: comparison with valproic acid autistic model. Pharmacol Biochem Behav 111, 102–110.

Bailey, K.R., Crawley J.N. (2009). Methods of Behavior Analysis in Neuroscience, 2nd edition, 2nd edn (Boca Raton (FL): http://www.crcpress.com/).

Basu, S.N., Kollu, R., and Banerjee-Basu, S. (2009). AutDB: a gene reference resource for autism research. Nucleic Acids Res 37, D832–836.

Bidinosti, M., Botta, P., Kruttner, S., Proenca, C.C., Stoehr, N., Bernhard, M., Fruh, I., Mueller, M., Bonenfant, D., Voshol, H., et al. (2016). CLK2 inhibition ameliorates autistic features associated with SHANK3 deficiency. Science 351, 1199–1203.

Bozdagi, O., Tavassoli, T., and Buxbaum, J.D. (2013). Insulin-like growth factor-1 rescues synaptic and motor deficits in a mouse model of autism and developmental delay. Mol Autism 4, 9.

Chadman, K.K., Guariglia, S.R., and Yoo, J.H. (2012). New directions in the treatment of autism spectrum disorders from animal model research. Expert Opin Drug Discov 7, 407–416.

Chen, S., Fragoza, R., Klei, L., Liu, Y., Wang, J., Roeder, K., Devlin, B., and Yu, H. (2018). An interactome perturbation framework prioritizes damaging missense mutations for developmental disorders. Nat Genet 50, 1032–1040.

Curwen, J.O., and Wedge, S.R. (2009). The use and refinement of rodent models in anti-cancer drug discovery: a review. Altern Lab Anim 37, 173–180.

Darnell, J.C., Van Driesche, S.J., Zhang, C., Hung, K.Y., Mele, A., Fraser, C.E., Stone, E.F., Chen, C., Fak, J.J., Chi, S.W., et al. (2011). FMRP stalls ribosomal translocation on mRNAs linked to synaptic function and autism. Cell 146, 247–261.

Duffney, L.J., Zhong, P., Wei, J., Matas, E., Cheng, J., Qin, L., Ma, K., Dietz, D.M., Kajiwara, Y., Buxbaum, J.D., et al. (2015). Autism-like Deficits in Shank3-Deficient Mice Are Rescued by Targeting Actin Regulators. Cell Rep 11, 1400–1413.

Foley, K.A., MacFabe, D.F., Kavaliers, M., and Ossenkopp, K.P. (2015). Sexually dimorphic effects of prenatal exposure to lipopolysaccharide, and prenatal and postnatal exposure to propionic acid, on acoustic startle response and prepulse inhibition in adolescent rats: relevance to autism spectrum disorders. Behav Brain Res 278, 244–256.

Harony-Nicolas, H., Kay, M., Hoffmann, J.D., Klein, M.E., Bozdagi-Gunal, O., Riad, M., Daskalakis, N.P., Sonar, S., Castillo, P.E., Hof, P.R., et al. (2017). Oxytocin improves behavioral and electrophysiological deficits in a novel Shank3-deficient rat. Elife 6.

Ji, X., Kember, R.L., Brown, C.D., and Bucan, M. (2016). Increased burden of deleterious variants in essential genes in autism spectrum disorder. Proc Natl Acad Sci U S A 113, 15054–15059.

Kalkbrenner, A.E., Windham, G.C., Serre, M.L., Akita, Y., Wang, X., Hoffman, K., Thayer, B.P., and Daniels, J.L. (2015). Particulate matter exposure, prenatal and postnatal windows of susceptibility, and autism spectrum disorders. Epidemiology 26, 30–42.

Kojima, M., Yassin, W., Owada, K., Aoki, Y., Kuwabara, H., Natsubori, T., Iwashiro, N., Gonoi, W., Takao, H., Kasai, K., et al. (2018). Neuroanatomical Correlates of Advanced Paternal and Maternal Age at Birth in Autism Spectrum Disorder. Cereb Cortex.

Krishnan, V., Stoppel, D.C., Nong, Y., Johnson, M.A., Nadler, M.J., Ozkaynak, E., Teng, B.L., Nagakura, I., Mohammad, F., Silva, M.A., et al. (2017). Autism gene Ube3a and seizures impair sociability by repressing VTA Cbln1. Nature 543, 507–512.

Kumar, A., Wadhawan, R., Swanwick, C.C., Kollu, R., Basu, S.N., and Banerjee-Basu, S. (2011). Animal model integration to AutDB, a genetic database for autism. BMC Med Genomics 4, 15.

Larrieu, T., Cherix, A., Duque, A., Rodrigues, J., Lei, H., Gruetter, R., and Sandi, C. (2017). Hierarchical Status Predicts Behavioral Vulnerability and Nucleus Accumbens Metabolic Profile Following Chronic Social Defeat Stress. Curr Biol 27, 2202–2210 e2204.

Laugeray, A., Herzine, A., Perche, O., Hebert, B., Aguillon-Naury, M., Richard, O., Menuet, A., Mazaud-Guittot, S., Lesne, L., Briault, S., et al. (2014). Pre- and postnatal exposure to low dose glufosinate ammonium induces autism-like phenotypes in mice. Front Behav Neurosci 8, 390.

Leblond, C.S., Nava, C., Polge, A., Gauthier, J., Huguet, G., Lumbroso, S., Giuliano, F., Stordeur, C., Depienne, C., Mouzat, K., et al. (2014). Meta-analysis of SHANK Mutations in Autism Spectrum Disorders: a gradient of severity in cognitive impairments. PLoS Genet 10, e1004580.

Li, K., Li, L., Cui, B., Gai, Z., Li, Q., Wang, S., Yan, J., Lin, B., Tian, L., Liu, H., et al. (2018). Early Postnatal Exposure to Airborne Fine Particulate Matter Induces Autism-like Phenotypes in Male Rats. Toxicol Sci 162, 189–199.

Manvich, D.F., Webster, K.A., Foster, S.L., Farrell, M.S., Ritchie, J.C., Porter, J.H., and Weinshenker, D. (2018). The DREADD agonist clozapine N-oxide (CNO) is reverse-metabolized to clozapine and produces clozapine-like interoceptive stimulus effects in rats and mice. Sci Rep 8, 3840.

Martinez-Cerdeno, V. (2017). Dendrite and spine modifications in autism and related neurodevelopmental disorders in patients and animal models. Dev Neurobiol 77, 393–404.

Mei, Y., Monteiro, P., Zhou, Y., Kim, J.A., Gao, X., Fu, Z., and Feng, G. (2016). Adult restoration of Shank3 expression rescues selective autistic-like phenotypes. Nature 530, 481–484.

Monteiro, P., and Feng, G. (2017). SHANK proteins: roles at the synapse and in autism spectrum disorder. Nat Rev Neurosci 18, 147–157.

Murugan, M., Jang, H.J., Park, M., Miller, E.M., Cox, J., Taliaferro, J.P., Parker, N.F., Bhave, V., Hur, H., Liang, Y., et al. (2017). Combined Social and Spatial Coding in a Descending Projection from the Prefrontal Cortex. Cell 171, 1663–1677 e1616.

O’Donnell, P. (2013). Of mice and men: what physiological correlates of cognitive deficits in a mouse model of schizophrenia tell us about psychiatric disease. Neuron 80, 265–266.

Rubenstein, J.L., and Merzenich, M.M. (2003). Model of autism: increased ratio of excitation/inhibition in key neural systems. Genes Brain Behav 2, 255–267.

Schulz, D., Huston, J.P., Buddenberg, T., and Topic, B. (2007). “Despair” induced by extinction trials in the water maze: relationship with measures of anxiety in aged and adult rats. Neurobiol Learn Mem 87, 309–323.

Slawinski, B.L., Talge, N., Ingersoll, B., Smith, A., Glazier, A., Kerver, J., Paneth, N., and Racicot, K. (2018). Maternal cytomegalovirus sero-positivity and autism symptoms in children. Am J Reprod Immunol 79, e12840.

Smith, V., and Brown, N. (2014). Prenatal valproate exposure and risk of autism spectrum disorders and childhood autism. Arch Dis Child Educ Pract Ed 99, 198.

Smithies, O. (1993). Animal models of human genetic diseases. Trends Genet 9, 112–116.

Speed, H.E., Kouser, M., Xuan, Z., Reimers, J.M., Ochoa, C.F., Gupta, N., Liu, S., and Powell, C.M. (2015). Autism-Associated Insertion Mutation (InsG) of Shank3 Exon 21 Causes Impaired Synaptic Transmission and Behavioral Deficits. J Neurosci 35, 9648–9665.

Victorino, D.B., Bederman, I.R., and Costa, A.C.S. (2017). Pharmacokinetic Properties of Memantine after a Single Intraperitoneal Administration and Multiple Oral Doses in Euploid Mice and in the Ts65Dn Mouse Model of Down’s Syndrome. Basic Clin Pharmacol Toxicol 121, 382–389.

von Bubnoff, A. (2008). Mighty mice. Scientists are still improving the humanized mouse model but are optimistic about its future role in evaluating AIDS vaccine candidates. IAVI Rep 12, 1, 8–11.

Wagner, G.C., Reuhl, K.R., Cheh, M., McRae, P., and Halladay, A.K. (2006). A new neurobehavioral model of autism in mice: pre- and postnatal exposure to sodium valproate. J Autism Dev Disord 36, 779–793.

Wang, X., Bey, A.L., Katz, B.M., Badea, A., Kim, N., David, L.K., Duffney, L.J., Kumar, S., Mague, S.D., Hulbert, S.W., et al. (2016). Altered mGluR5-Homer scaffolds and corticostriatal connectivity in a Shank3 complete knockout model of autism. Nat Commun 7, 11459.

Wang, X., Xu, Q., Bey, A.L., Lee, Y., and Jiang, Y.H. (2014). Transcriptional and functional complexity of Shank3 provides a molecular framework to understand the phenotypic heterogeneity of SHANK3 causing autism and Shank3 mutant mice. Mol Autism 5, 30.

Washington, M.K., Powell, A.E., Sullivan, R., Sundberg, J.P., Wright, N., Coffey, R.J., and Dove, W.F. (2013). Pathology of rodent models of intestinal cancer: progress report and recommendations. Gastroenterology 144, 705–717.

Wickham, H. (2014). Tidy data. Journal of Statistical Software 59, 1–23.

Wickham, H. (2016). Ggplot2 (New York, NY: Springer Science+Business Media, LLC).

Won, H., Lee, H.R., Gee, H.Y., Mah, W., Kim, J.I., Lee, J., Ha, S., Chung, C., Jung, E.S., Cho, Y.S., et al. (2012). Autistic-like social behaviour in Shank2-mutant mice improved by restoring NMDA receptor function. Nature 486, 261–265.

Zeng, X.S., Geng, W.S., and Jia, J.J. (2018). Neurotoxin-Induced Animal Models of Parkinson Disease: Pathogenic Mechanism and Assessment. ASN Neuro 10, 1759091418777438.

Zhou, Y., Kaiser, T., Monteiro, P., Zhang, X., Van der Goes, M.S., Wang, D., Barak, B., Zeng, M., Li, C., Lu, C., et al. (2016). Mice with Shank3 Mutations Associated with ASD and Schizophrenia Display Both Shared and Distinct Defects. Neuron 89, 147–162.

